# Inflammasome-independent role for NLRP3 in controlling innate anti-helminth immunity and tissue repair in the lung

**DOI:** 10.1101/606392

**Authors:** AL Chenery, R Alhallaf, Z Agha, J Ajendra, JE Parkinson, MM Cooper, BHK Chan, RM Eichenberger, LA Dent, AAB Robertson, A Kupz, D Brough, A Loukas, TE Sutherland, JE Allen, PR Giacomin

## Abstract

Alternatively activated macrophages are essential effector cells during type 2 immunity and tissue repair following helminth infections. We previously showed that Ym1, an alternative activation marker, can drive innate IL-1R-dependent neutrophil recruitment during infection with the lung-migrating nematode, *Nippostrongylus brasiliensis* suggesting a potential role for the inflammasome in the IL-1-mediated innate response to infection. While inflammasome proteins such as NLRP3 have important pro-inflammatory functions in macrophages, their role during type 2 responses and repair are less defined. We therefore infected *Nlrp3*^−/−^ mice with *N. brasiliensis*. Unexpectedly, compared to WT mice, infected *Nlrp3*^−/−^ mice had increased neutrophilia and eosinophilia, correlating with enhanced worm killing but at the expense of increased tissue damage and delayed lung repair. Transcriptional profiling showed that infected *Nlrp3*^−/−^ mice exhibited elevated type 2 gene expression compared to WT mice. Notably, inflammasome activation was not evident early post-infection with *N. brasiliensis* and in contrast to *Nlrp3*^−/−^ mice, anti-helminth responses were unaffected in caspase-1/11 deficient or WT mice treated with the NLRP3-specific inhibitor MCC950. Together these data suggest that NLRP3 can constrain lung neutrophilia and helminth killing and negatively regulate type 2 immune responses in an inflammasome-independent manner.

## Introduction

The lungs are a vital barrier organ that must respond appropriately to both pathogens and innocuous antigens to maintain homeostasis. After tissue injury or an infectious insult, the lungs rapidly initiate immune resolution and repair mechanisms to maintain their essential physiological function. Alveolar macrophages (Mφs) help maintain homeostasis and can drive either pro-inflammatory or pro-resolution responses in the lung tissue microenvironment, depending on the stimulus. Classically activated Mφs upregulate anti-microbial factors such as NO, TNFα, IL-6, and IL-1β in response to pathogen or damage-associated molecular patterns (PAMPs or DAMPs respectively) (1). Conversely, during helminth infections, Mφs can become alternatively activated in response to IL-4Rα signalling, upregulating arginase, resistin-like molecule (RELM)-α, chitinase-like protein Ym1, and specific matrix metalloproteinases (MMPs), all important type 2 effector molecules with wound healing functions (2–4). IL-4Rα-activated Mφs are known to mediate tissue repair in the skin (4) but also control airway inflammation and haemorrhage following the acute lung injury caused by primary infection with lung migrating nematodes (5). Upon secondary infection, IL-4Rα-activated Mφs act in cooperation with anti-parasitic neutrophil, eosinophil, ILC2 and memory Th2 cell responses to mediate parasite control (6–9).

Mφ-derived Ym1 is a key feature of acute lung injury following infection with the nematode *Nippostrongylus brasiliensis*, with further increases in expression in response to the subsequent Th2 cell response (2). However, even in the absence of IL-4 under steady state conditions, alveolar Mφs express substantial amounts of Ym1 (2). We have previously shown that as early as 2 days post-*N. brasiliensis* infection, Ym1 has played a prominent role leading to recruitment of neutrophils that swarm around larvae in the lungs and promote worm killing (10). IL-17A-producing γδT cells are key to Ym1-mediated neutrophilia. Ectopic overexpression of Ym1 drives *Il1b* expression whilst neutralising Ym1 prevents expression of *Il1b* following *N. brasiliensis* infection (10). Because blockade of the IL-1 receptor reduces the ability of Ym1 to induce IL-17-producing γδT cells and associated neutrophilia (10) we and others (11) hypothesized that Ym1 may activate an inflammasome, leading to IL-1 release. In addition, Ym1 is well known to crystallize under conditions of chronic inflammation (12) and crystalline/particulate material can activate the NLRP3 (NLR family member NACHT, LRR and PYD domains-containing protein 3) inflammasome suggesting that Ym1 may be a specific activator of NLRP3. Therefore, we predicted that NLRP3 inflammasome activation by alveolar Mφs would drive early neutrophilic responses important for lung-stage larval killing during *N. brasiliensis* infection. However, questions still remain about how tissue signals, like inflammasome activation, may influence alveolar Mφ activation during the course of lung injury and repair.

While the NLRP3 inflammasome has been widely studied in the context of classically activated Mφs, comparatively little is known about whether NLRP3 plays a role during alternative activation and type 2 inflammatory settings in which Ym1 is highly induced. Our previous studies show that NLRP3 constrains type 2 responses during infection with the gut-dwelling helminth parasite *Trichuris muris*, with a major effect on the development of adaptive Th2 cell immunity (13). In the present study, we addressed whether the NLRP3 inflammasome contributed to acute lung injury, innate immune cell activation and lung repair during infection with *N. brasiliensis*. Counter to expectations, we discovered that NLRP3 limited rather than promoted early lung neutrophilia. Additionally, NLRP3 impaired host anti-helminth effector mechanisms and other type 2 immune responses, but may have been important for limiting infection-induced tissue damage.

## Results

### NLRP3 restricts early innate immune cell recruitment to the lung

To test our hypothesis that NLRP3 is required for the early recruitment of neutrophils into the lung during infection with lung-migrating helminths, WT and *Nlrp3*^−/−^ mice were infected with *N. brasiliensis*. Lung tissue cells were isolated and analysed on day 2 post-infection, where there was a peak of injury and neutrophilia. As expected, *N. brasiliensis* infection of WT mice led to increased frequencies (**Figure 1a**) and total numbers (**Figure 1b**) of eosinophils and neutrophils in the lung, compared to naïve mice. Unexpectedly, infected *Nlrp3*^−/−^ mice had increased numbers of recruited neutrophils and eosinophils compared to infected WT mice. Total numbers of alveolar Mφs were not significantly different between all groups (**Figure 1a-b**). Cellular content of bronchoalveolar lavage (BAL) fluid was not analysed because all infected samples on day 2 were bloody due to lung damage. Together, these data suggest that rather than promote granulocyte recruitment to the lung, NLRP3 may instead limit the recruitment of neutrophils and eosinophils.

**Figure 1.**
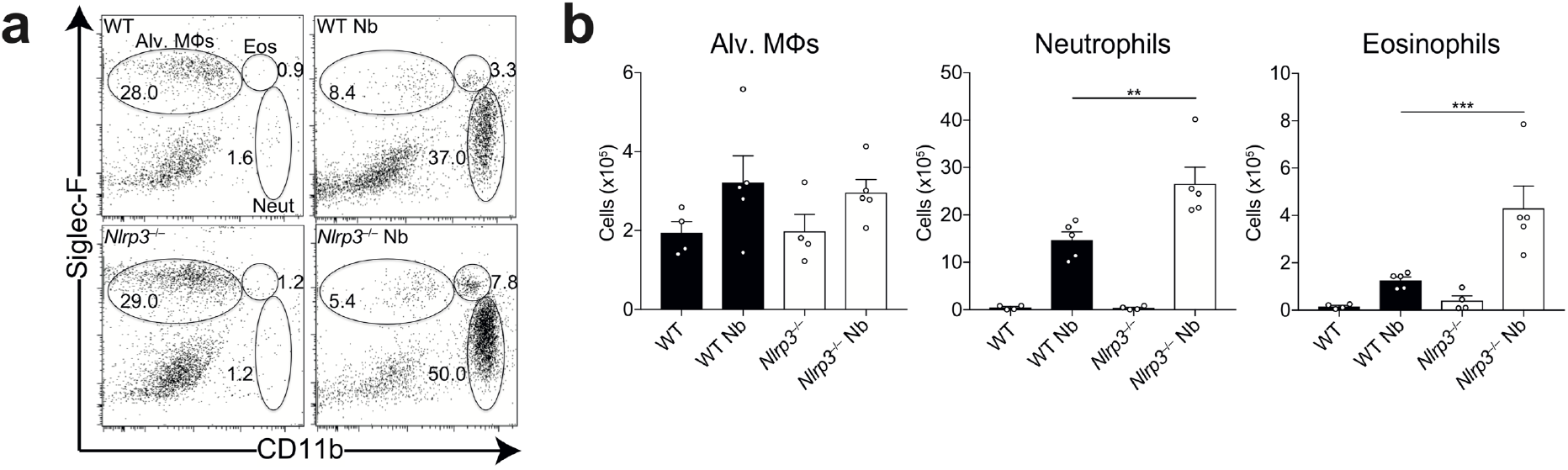
NLRP3 constrains lung innate cell recruitment during *N. brasiliensis* infection. WT and *Nlrp3*^−/−^ mice were infected with *N. brasiliensis* (Nb) and day 2 post-infected lung alveolar (alv.) Mφ (CD11b^lo^Siglec-F^+^CD11c^+^), neutrophil (CD11b^+^Siglec-F^−^Ly6G^+^) and eosinophil (CD11b^+^Siglec-F^+^) (**a**) frequencies of live cells and (**b**) absolute numbers were measured by flow cytometry. Data are representative (mean ± s.e.m.) of 4 individual experiments with 3-5 mice per group (per experiment). \*\**P*<0.01, \*\*\**P*<0.001, (one-way ANOVA and Tukey-Kramer *post hoc* test).

### NLRP3 limits innate anti-helminth immunity in the lung

We next addressed whether the increased granulocyte infiltration in infected *Nlrp3*^−/−^ mice correlated with increased anti-helminth immunity. Rapid neutrophil recruitment to the lung in particular is known to be important for immunity to *N. brasiliensis* (10, 14). Thus, we assessed numbers of larvae in WT and *Nlrp3*^−/−^ mice during primary *N. brasiliensis* infection. Consistent with the enhanced early granulocyte response (**Figure 1b**), *Nlrp3*^−/−^ mice displayed reduced numbers of lung-stage L4 larvae at both day 1 and day 2 post-infection compared to WT mice (**Figure 2a**). Consequently, *Nlrp3*^−/−^ mice also displayed less parasite migration to the intestine at days 4 and day 6 post-infection (**Figure 2b**) and a decrease in shed fecal eggs (**Figure 2c**). *Nlrp3*^−/−^ mice also displayed increased presence of goblet cells in the small intestine (**Figure 2d**), consistent with an increased effector type 2 response (15). Together, these data suggest that NLRP3 may constrain protective innate immune responses in the lung following *N. brasiliensis* infection, potentially by suppressing neutrophil and eosinophil accumulation.

**Figure 2.**
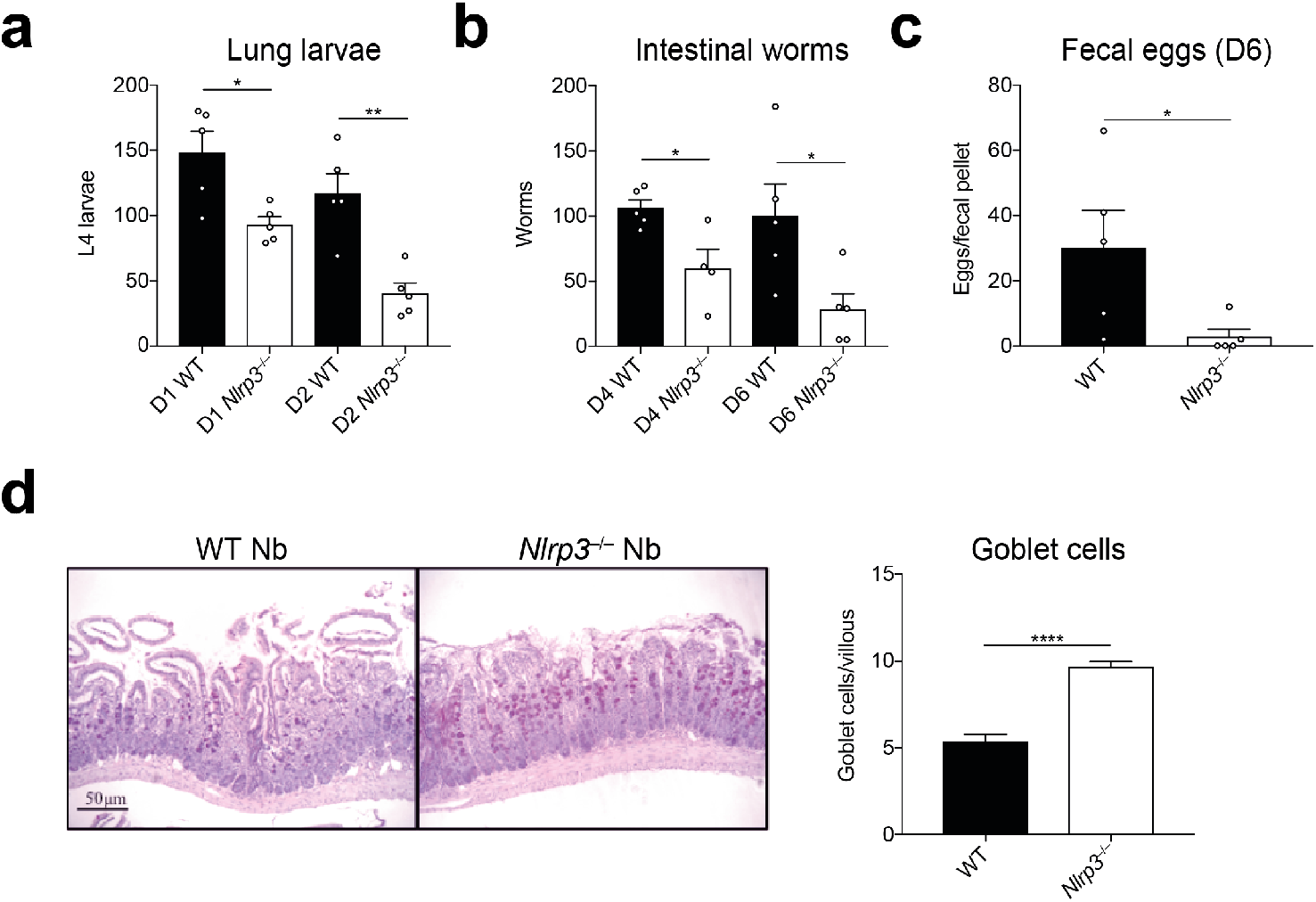
Restriction of anti-helminth responses by NLRP3. WT and *Nlrp3*^−/−^ mice were infected with 500 *N. brasiliensis* (Nb) L3s and (**a**) lung stage L4 larvae were quantified on days 1 and 2 post-infection. (**b**) Adult worms in the small intestine (days 4 and 6 post-infection) and (**c**) fecal parasite eggs shed in the stool (day 6) were quantified (per fecal pellet). (**d**) Representative images of proximal small intestine from day 6 infected mice, stained with Alcian Blue/Periodic Acid-Schiff. Goblet cells were enumerated from 10 randomly selected villi per mouse. Data from **a-d** are representative (mean ± s.e.m.) of 3 individual experiments with 3-5 mice per group (per experiment). \**P*<0.05, \*\**P*<0.01 (one-way ANOVA and Tukey-Kramer *post hoc* test or unpaired two-tailed student’s *t* test).

### NLRP3 regulates lung repair after helminth infection

The data thus far demonstrated that *Nlrp3*^−/−^ mice displayed elevated neutrophilia and eosinophilia in the lung and protective immunity in the early stages of primary infection, accompanied by an enhanced type 2 effector response in the intestine. Both types of granulocytes may contribute to lung tissue damage during *N. brasiliensis* infection and allergic responses (5, 10) but type 2 responses are typically tissue protective (16). It was therefore important to assess whether elevated granulocytic responses in *Nlrp3*^−/−^ mice impacted the resolution of inflammation and repair of lung tissue damage during the later stage of the infection model (17, 18). To assess airway cell infiltration, BAL was performed on mice at day 7 post-*N. brasiliensis* infection, at a timepoint when haemorrhaging has normally ceased and larvae have completely exited the lung tissue. As expected, eosinophils were the predominant cell type elevated in the airways of infected WT mice compared to naïve controls along with increased alveolar Mφs and neutrophils (**Figure 3a**). Critically, all three cell types were significantly elevated in infected *Nlrp3*^−/−^ mice compared to infected WT mice, suggesting a persistence of cellular inflammation in the airways. This was consistent with evidence of increased lung tissue pathology in *Nlrp3*^−/−^ mice compared to WT mice after infection (**Figure 3b-c**). Specifically, while lungs from infected WT mice displayed modest airway cell infiltration and features of successful tissue repair, *Nlrp3*^−/−^ lungs had more prominent and dispersed immune cell foci and larger gaps in the alveoli indicating impaired repair (**Figure 3c**). To quantify lung damage, we performed fractal/lacunarity analysis of entire lung lobes which has previously been shown to robustly measure lung pathology, correlating well with traditional stereological measurements such as mean linear intercept (19). Lacunarity (Λ), a measure of heterogeneity/gaps in a structure that correlates with airway damage, was increased for infected *Nlrp3*^−/−^ lungs compared to infected WT lungs which showed no significant differences over WT naïve controls (**Figure 3b**). We did not detect differences in tissue pathology between WT and *Nlrp3*^−/−^ lungs as early as day 2 post-infection (**Figure S1**). Together, these data reveal that NLRP3 is required for the resolution of inflammation and the timely initiation of repair processes in the lung following injury by helminth infection.

**Figure 3.**
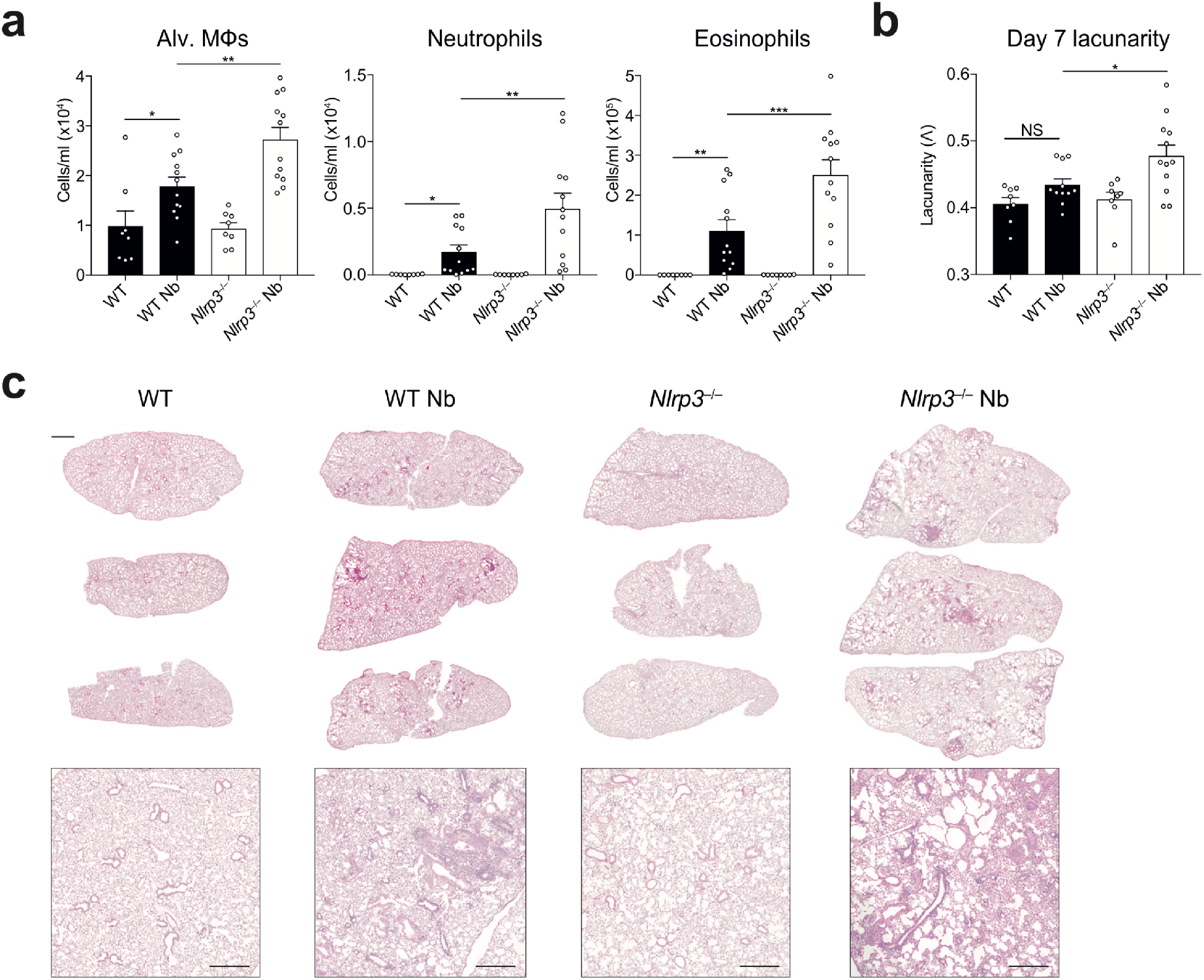
NLRP3 regulates lung tissue repair following *N. brasiliensis* infection. WT and *Nlrp3*^−/−^ mice were infected with *N. brasiliensis* (Nb) and (**a**) day 7 post-infected BAL alveolar (alv.) Mφ (CD11b^−^Siglec-F^+^CD11c^+^), neutrophil (CD11b^+^Siglec-F^−^Ly6G^+^) and eosinophil (CD11b^+^Siglec-F^+^) absolute numbers were measured by flow cytometry. Haematoxylin/eosin staining was performed on lung sections and imaged followed by (**b**) quantification of lacunarity (Λ) to assess lung damage on day 7. (**c**) Representative image insets are shown on top with low magnification scans of entire left lung lobes from individual mice shown below (top panel scale bar = 1000 μm, bottom panel scale bar = 500 μm). Data were pooled (**a-b**; mean ± s.e.m.) from 3 individual experiments with 3-5 mice per group (per experiment). \**P*<0.05, \*\**P*<0.01, \*\*\**P*<0.001 (one-way ANOVA and Tukey-Kramer *post hoc* test).

### NLRP3 constrains type 2 cytokine responses and expression of Ym1

The failure of *Nlrp3*^−/−^ mice to repair as well as WT mice was not fully consistent with their enhanced eosinophilia and goblets cells, which would suggest that NLRP3 reduces type 2 immunity. Lung tissue repair following *N. brasiliensis* infection is dependent on type 2 responses including the expansion of IL-4Rα-activated Mφs (5). Critically, while innate sources of Ym1 induce neutrophilia (10), during the later reparative stage, IL-4Rα-induced Ym1 has a direct role in repair (2). We therefore examined whether type 2 responses are dysregulated in *Nlrp3*^−/−^ mice during *N. brasiliensis* infection. Despite a failure to repair, compared to infected WT mice, infected *Nlrp3*^−/−^ mice exhibited enhanced IL-4 protein levels in the lungs as early as day 2 post-infection (**Figure 4a**). By day 7 post-infection, using qRT-PCR we found that *Chil3* (Ym1) expression was significantly increased in the lungs of *Nlrp3*^−/−^ mice compared to WT mice (**Figure 4b**). Similarly, there were significant increases in secreted Ym1 protein levels in the BAL fluid of infected *Nlrp3*^−/−^ mice (**Figure 4c**). To define the cellular source of Ym1, we performed intracellular cytokine staining and flow cytometry focussing on alveolar Mφs and neutrophils that are known sources of Ym1 during *N. brasiliensis* infection (2). We found that infection caused a drop in the geometric mean fluorescence intensity (gMFI) of Ym1 staining in both alveolar Mφs and neutrophils (**Figure 4d)**, consistent with release of Ym1 into the airways, as observed in the BAL fluid (**Figure 4c**). However, *Nlrp3*^−/−^ alveolar Mφs had a greater drop in gMFI for Ym1 following infection compared to WT alveolar Mφs. Conversely, Ym1 release by neutrophils was equivalent in infected WT and *Nlrp3*^−/−^ mice. Therefore, NLRP3 constrains expression of IL-4 and Ym1 in the lung during infection with *N. brasiliensis* and may control release of Ym1 by alveolar Mφs in the airways.

**Figure 4.**
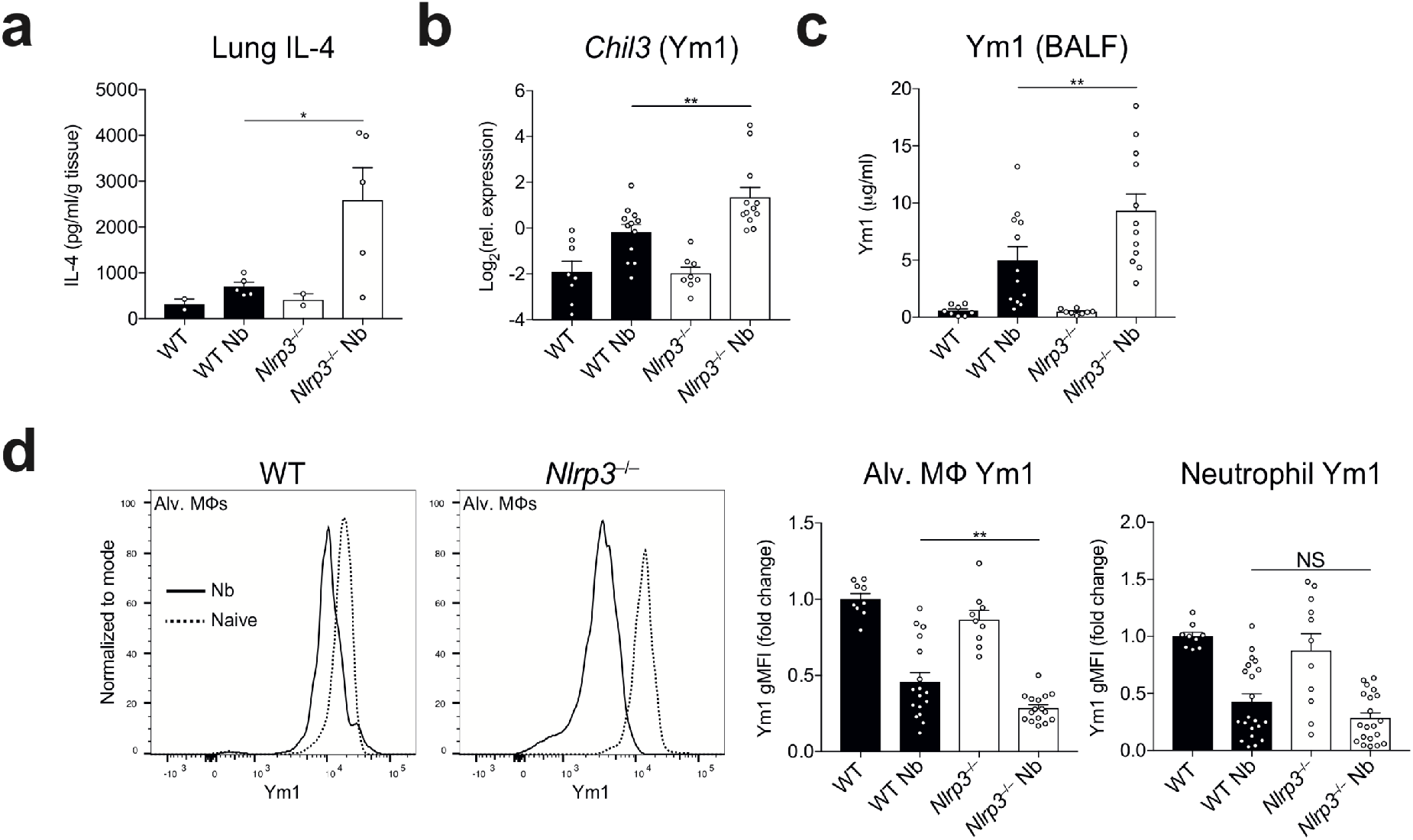
NLRP3 regulates Ym1 following infection with *N. brasiliensis*. After infection of WT and *Nlrp3*^−/−^ mice with *N. brasiliensis* (Nb), (**a**) IL-4 protein levels were measured in lung tissues by ELISA on day 2 post-infection (normalised to tissue weight). (**b**) *Chil3* (Ym1) gene expression was measured by qRT-PCR on day 7 post-infection (log_2_ expression relative to *Rpl13a*). (**c**) BALF Ym1 was measured by ELISA and (**d**) intracellular expression by geometric mean fluorescence intensity (gMFI) (fold change over WT control) by alveolar (alv.) Mφs and neutrophils were quantified by flow cytometry on day 7 post-infection. Data are representative (**a**; mean ± s.e.m.) or pooled (**b-d**; mean ± s.e.m.) from 3 individual experiments with 3-5 mice per group (per experiment). \**P*<0.05, \*\**P*<0.01, (one-way ANOVA and Tukey-Kramer *post hoc* test).

### Type 2 gene expression is regulated by NLRP3

To more broadly characterise the immune processes potentially dysregulated in *Nlrp3*^−/−^ mice, we performed differential gene expression analysis of whole lung RNA on day 7 post-infection with *N. brasiliensis* using the NanoString nCounter® Myeloid Innate Immunity panel. We observed differential expression of well-characterised genes associated with the later stages of *N. brasiliensis* infection such as *Rnase2a*, *Retnla* and *Chil3/4* that were, as expected, highly upregulated in infected WT mice relative to uninfected mice (**Figure S3**). Under basal conditions, few genes were found to be downregulated in *Nlrp3*^−/−^ mice compared to naïve WT mice (**Figure S2**). However, in *N. brasiliensis*-infected mice, 84 genes were differentially expressed between *Nlrp3*^−/−^ mice compared to WT (*P* < 0.05) (**Figure 5a-b**). Although no genes remained statistically significant after Benjamini-Hochburg multiple test correction (20), principle components analysis (PCA) and hierarchical clustering (**Figures 5c-d**) showed clear separation of gene expression patterns between *N. brasiliensis*-infected WT and *Nlrp3*^−/−^ mice. Increased sample numbers were used to validate specific genes by qRT-PCR. Infected *Nlrp3*^−/−^ mice had significantly higher expression of the type 2 markers *Arg1* and *Ccl24* as well as an increase in the neutrophil-associated *Cxcl3* compared to infected WT mice (**Figure 5e**). Despite not reaching significance in the NanoString analysis, qRT-PCR analysis showed that *Il4* expression was found to be significantly increased in infected *Nlrp3*^−/−^ mice compared to infected WT mice, confirming the protein levels observed in **Figure 4a**. Additional genes whose expression was increased in infected *Nlrp3*^−/−^ mice included the matrix metalloprotease-encoding *Mmp13* and the alveolar Mφ-expressed gene *Ear6* (21). Thus, NLRP3 appears to negatively regulate type 2 gene expression in the lung following tissue injury caused by helminth infection and may control neutrophil chemotactic recruitment and alveolar Mφ function.

**Figure 5.**
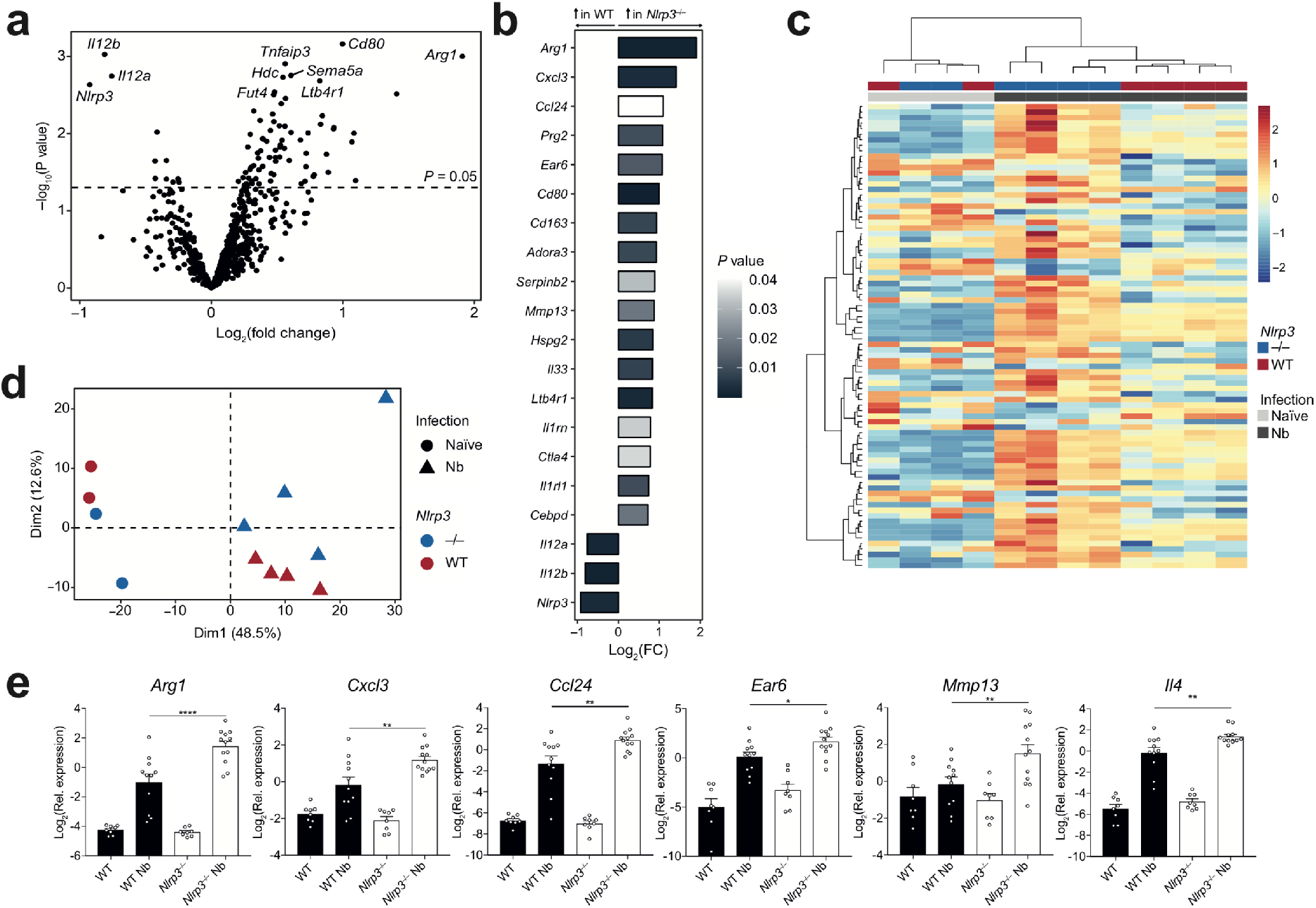
NLRP3 regulates type 2 gene expression in the lung following *N. brasiliensis* infection. Whole lung RNA from WT and *Nlrp3*^−/−^ mice on day 7 post-infection with *N. brasiliensis* (Nb) was analysed by NanoString. (**a**) Volcano plot showing differentially expressed genes between Nb-infected WT and *Nlrp3*^−/−^ mice. (**b**) Top 20 differentially regulated genes between Nb-infected WT and *Nlrp3*^−/−^ mice (bar colour indicates statistical significance). (**c**) Unsupervised, hierarchically clustered heatmap of genes differentially expressed 7 days post Nb-infection in WT and *Nlrp3*^−/−^ mice. (**d**) Principle components analysis (PCA) of naïve and Nb-infected WT and *Nlrp3*^−/−^ mice. (**e**) Candidate differentially expressed genes validated by qRT-PCR (log_2_ expression relative to *Rpl13a*). Data were from a single Nanostring run (**a-d**; n=2-4 mice per group) or pooled (**e**; mean ± s.e.m.) from 3 individual experiments with 3-5 mice per group (per experiment). \**P*<0.05, \*\**P*<0.01, \*\*\**P*<0.001, \*\*\*\**P*<0.0001 (one-way ANOVA and Tukey-Kramer *post hoc* test).

### Tissue resident Mφ alternative activation is regulated by NLRP3

Our finding that expression of *Chil3* (Ym1) was higher in *N. brasiliensis*-infected *Nlrp3*^−/−^ mice (**Figure 4**) is likely due to the enhanced quantities of IL-4 observed as several IL-4-responsive genes were observed to be upregulated in the absence of NLRP3 (**Figure 5**). However, it is also possible that NLRP3 enhances responsiveness of Mφs to IL-4Rα signalling. To test this, we utilized IL-4-complex (IL-4c) delivery into the peritoneal cavity which robustly induces expression of Ym1 and RELM-α in peritoneal Mφs (22). WT and *Nlrp3*^−/−^ mice were injected with IL-4c i.p. following which we analysed the CD11b^+^ myeloid compartment containing F4/80^lo^ monocyte-derived Mφs and F4/80^hi^Tim4^+^ tissue resident Mφs after 24 hours (**Figure 6a**). There would be no expectation that IL-4c delivery would induce the NLRP3 inflammasome and indeed we were unable to detect any evidence for inflammasome activation in the peritoneal Mφs (data not shown). As expected, both Mφ populations showed increased intracellular expression of Ym1 and RELM-α in WT mice after IL-4c injection. IL-4c-stimulated *Nlrp3*^−/−^ resident (F4/80^hi^Tim4^+^) Mφs had greater frequencies of both Ym1^+^ and RELM-α^+^ cells as well as an increased Ym1 gMFI compared to IL-4c-stimulated WT resident Mφs (**Figure 6b**). In contrast, we did not see enhanced expression of Ym1 or RELM-α above WT levels in *Nlrp3*^−/−^ monocyte-derived (F4/80^lo^) Mφs after IL-4c stimulation (**Figure 6c**). Similarly, *in vitro* IL-4-stimulated BMDMs from *Nlrp3*^−/−^ mice did not show increased alternative activation markers compared to WT BMDMs (**Figure S3**). Thus, it appears that *in vivo* NLRP3 can influence the Mφ response to IL-4, independent of the inflammasome, but these effects may be restricted to specific Mφs subpopulations, such as tissue resident rather than monocyte-derived Mφs.

**Figure 6.**
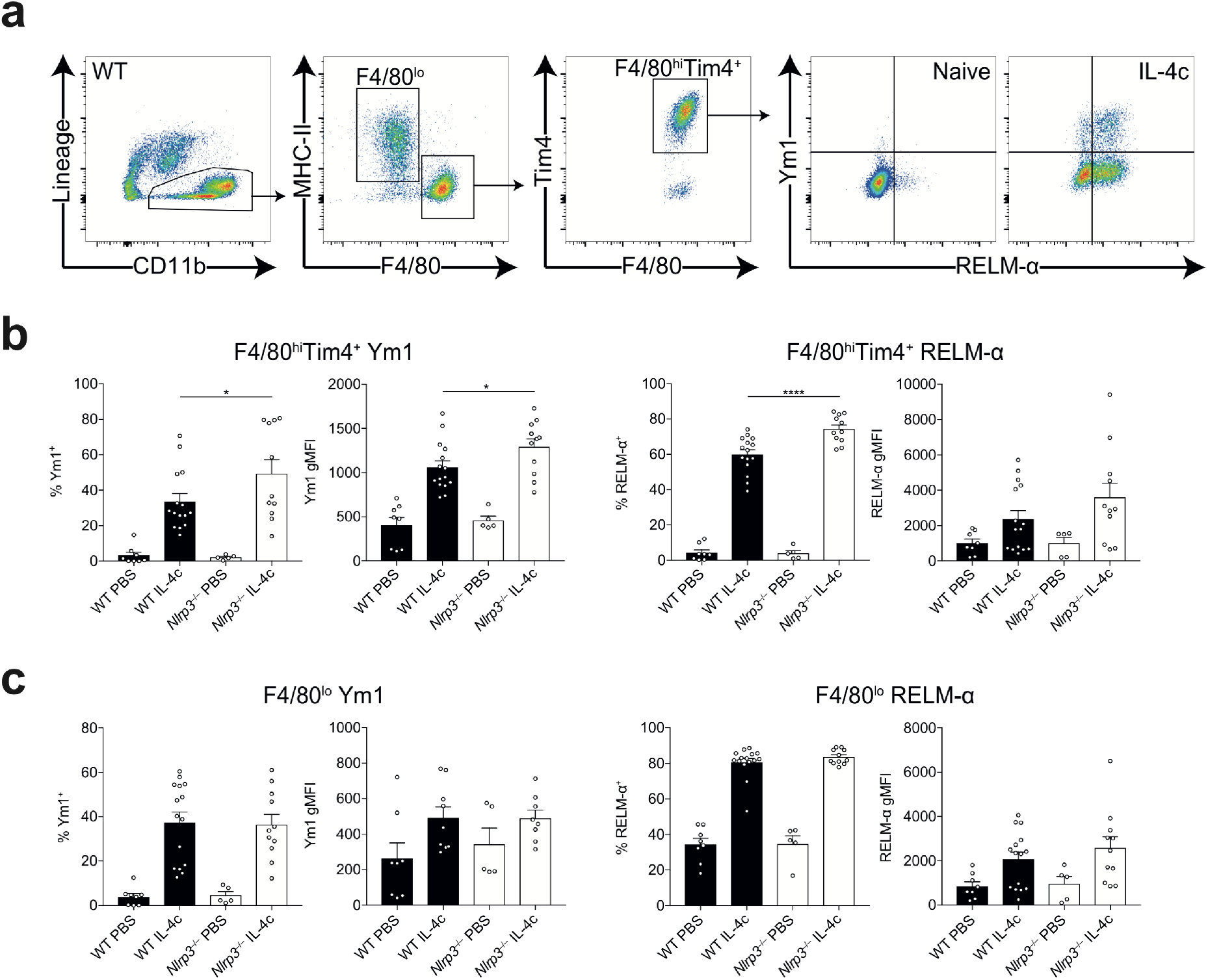
NLRP3 regulates resident Mφ alternative activation by IL-4. WT and *Nlrp3*^−/−^ mice were injected with either PBS or IL-4 complex (IL-4c) and (**a**) peritoneal CD11b^+^ cells containing F4/80^lo^ and F4/80^hi^Tim4^+^ Mφs where analysed by flow cytometry after 24 hours. Frequency and geometric mean fluorescence intensity (gMFI) of intracellular Ym1 and RELM-α were determined in (**b**) resident F4/80^hi^Tim4^+^ Mφs and (**c**) monocyte-derived F4/80^lo^ Mφs. Data are representative (**b** left panel; mean ± s.e.m.) or pooled (**b** right panel and **c**; mean ± s.e.m.) from 3 individual experiments with 3-5 mice per group (per experiment). \**P*<0.05, \*\*\*\**P*<0.0001 (one-way ANOVA and Tukey-Kramer *post hoc* test).

### Potential inflammasome-independent role of NLRP3 during lung anti-helminth responses

Thus far we have demonstrated that although NLRP3 suppressed type 2 immune responses, it also limited persistent lung inflammation after resolution of helminth infection. Our observation that NLRP3 constrained the early neutrophilic response was contrary to the expectation that infection with *N. brasiliensis* would result in inflammasome activation. Although NLRP3 is most commonly associated with forming an inflammasome, there are some reports of inflammasome-independent roles for NLRP3 (23, 24). Therefore, we investigated whether NLRP3 played inflammasome-dependent or -independent functions during *N. brasiliensis* infection. First, we examined expression of infection-induced production of pro-IL-1β within alveolar Mφs, a precursor for inflammasome activation. *N. brasiliensis* infection caused increased levels of pro-IL-1β within alveolar Mφs, however this infection-induced response was unaffected by NLRP3 deficiency (**Figure 7a**). Further, levels of mature IL-1β released into the airways (an indicator of inflammasome activation), were nearly undetectable in the BAL fluid in both WT and *Nlrp3*^−/−^ mice (**Figure 7b**), suggesting that inflammasome activation in the lung is not a major feature of *N. brasiliensis* infection. To further delineate whether inflammasome activation occurred following *N. brasiliensis* infection, we assessed the protease caspase-1. The inflammasome recruits and activates caspase-1 (25), which subsequently cleaves pro-IL-1β into its mature form (26). Activated caspase-1 was quantified *ex vivo* using a FAM-YVAD-FMK fluorochrome inhibitor of caspases (FLICA) probe on BAL cells on day 2 post-infection. Critically, there was no evidence of infection-induced increases in caspase-1 activity in alveolar Mφs (**Figure 7c**). Similarly, we observed no differences in caspase-1 activity within neutrophils and eosinophils between infected WT and *Nlrp3*^−/−^ mice.

**Figure 7.**
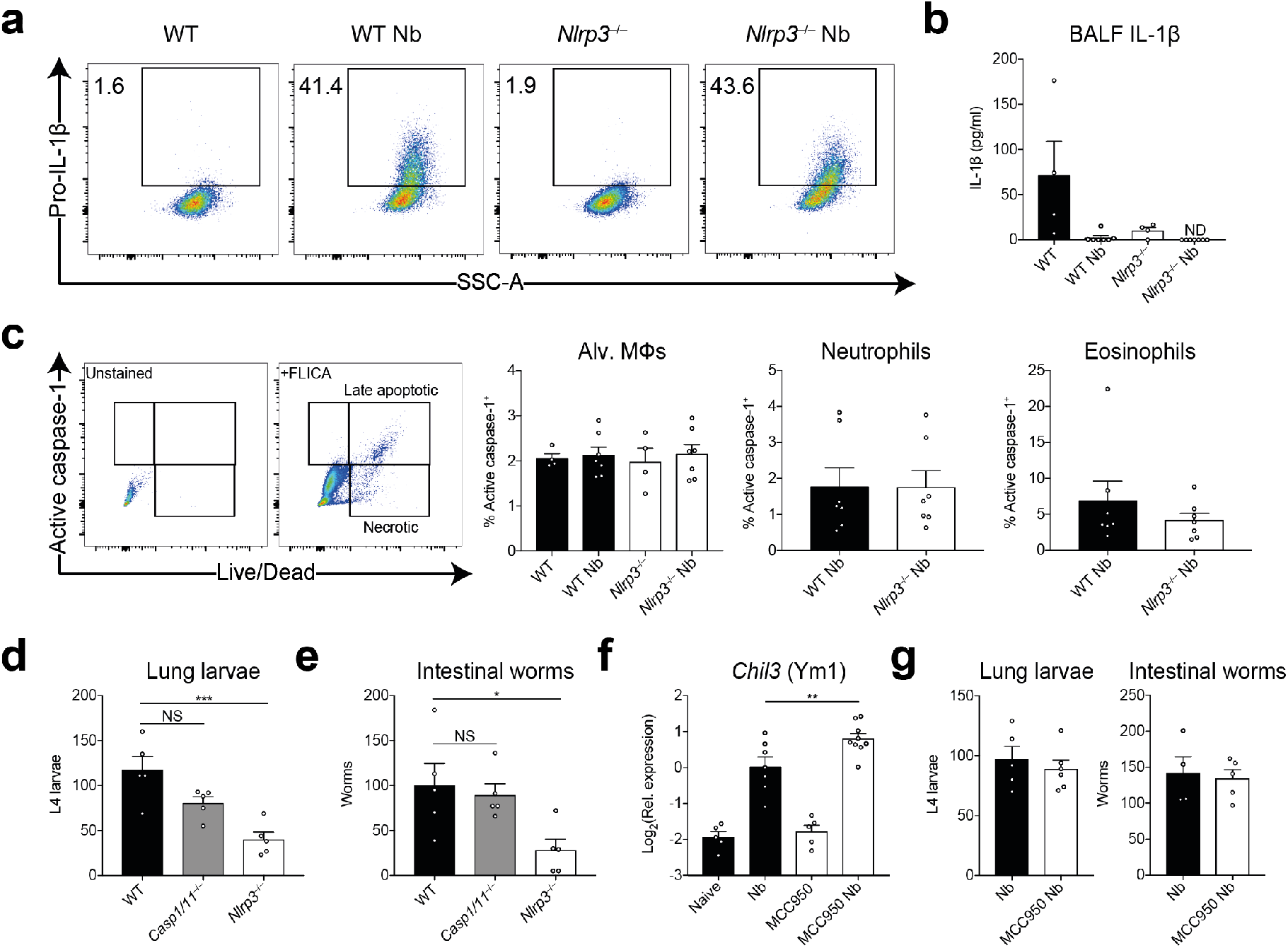
Inflammasome-independent role for NLRP3 during innate anti-helminth responses in the lung. WT and *Nlrp3*^−/−^ mice were infected with *N. brasiliensis* (Nb) and on day 2 post-infection (**a**) intracellular pro-IL-1β levels were measured in alveolar (Alv.) Mφs by flow cytometry, (**b**) released IL-1β in the BALF was quantified by ELISA and (**c**) frequency of active caspase-1 was measured *ex vivo* in BAL alveolar Mφs, neutrophils, and eosinophils by FAM-FLICA fluorescence. (**d**) On day 2 post-infection, L4 lung stage larvae and (**e**) day 6 adult intestinal worms were counted in WT, *Casp1/11*^−/−^ and *Nlrp3*^−/−^ mice. WT mice were treated with MCC950 during Nb infection and (**f**) *Chil3* (Ym1) expression in the lung was measured by qRT-PCR. (**g**) Lung larval burdens (day 2) and intestinal (day 6) worm burdens were counted following MCC950 treatment and Nb infection. Data are representative (**a-e, g**; mean ± s.e.m.) or pooled (**f**; mean ± s.e.m.) from 2 or 3 individual experiments with 3-5 mice per group (per experiment). \**P*<0.05, \*\**P*<0.01, \*\*\**P*<0.001 (one-way ANOVA and Tukey-Kramer *post hoc* test).

To more definitively establish whether inflammasome activation contributed to the observed phenotype of *Nlrp3*^−/−^ mice, we compared the response to *N. brasiliensis* infection in *Nlrp3*^−/−^ mice to that of *Casp1/11*^−/−^ mice which cannot mount either canonical or non-canonical NLRP3 inflammasome responses (27). While *Casp1/11*^−/−^ mice showed a trend towards increased immunity to lung-stage *N. brasiliensis* infection compared to WT mice, this was not statistically significant (**Figure 7d**) and adult worm burdens were equivalent to WT mice (**Figure 7e**). In contrast, *Nlrp3*^−/−^ mice displayed increased anti-parasitic immunity at the lung stage, characterised by significantly reduced lung larval numbers and adult worm burdens. In a final effort to establish a role for *N. brasiliensis*-induced inflammasome activation we treated WT mice with MCC950, an NLRP3-specific inflammasome small molecule inhibitor (28) throughout the course of infection (**Figure S5**). We failed to see changes in myeloid cell numbers in the BAL throughout the infection following MCC950 treatment (data not shown). By day 5 post-infection, we could see that mice treated with MCC950 had a reduction in BAL neutrophil numbers but had no change in alveolar Mφ or eosinophil numbers (**Figure S4**). However, we did observe increased expression of *Chil3* in the lungs of infected mice treated with MCC950 (**Figure 7f**). Although consistent with our findings in infected *Nlrp3*^−/−^ mice, other type 2 markers including IL-4 itself were unchanged in MCC950 treated or *Casp1/11*^−/−^ mice (data not shown). Further, in contrast to NLRP3 deficiency, there was no difference in lung-stage larvae numbers or intestinal worm burden between untreated and MCC950-treated mice (**Figure 7g**). Together, these data strongly suggest that NLRP3 plays an inflammasome-independent role in constraining neutrophil and anti-helminth immunity in the lung.

## Discussion

Whilst the role of NLRP3 in classical inflammatory settings has been well-characterised, the function of NLRP3 during type 2 inflammation has been more contentious. An early controversy focussed on whether the pro-type 2 adjuvant activity of alum is dependent on its ability to activate the NLRP3 inflammasome (29, 30). Subsequently, a variety of studies have found that NLRP3 either promotes (31–33) or restricts (27, 34) allergic responses and Th2 cells. Most recently, Perrson *et al.* demonstrated a profound ability of Charcot-Leyden crystals to promote type 2 immunity but the ability of the crystals to activate the NLRP3 inflammasome was not required for the adjuvant effect (35). Thus, the role of NLRP3 in type 2 immunity remains largely unresolved.

To help resolve this question, we have focussed on helminth infections, in which type 2 immunity is central for host protection. We previously found that the NLRP3 inflammasome played a major role in suppressing the adaptive type 2 immune response to intestinal helminth infection (in the caecum) with *T. muris* (13). In that study, MCC950 treatment and Caspase-1/11 deficiency was able to phenocopy *Nlrp3*^−/−^ mice during infection. In contrast, the present study showed that during the innate immune phase of *N. brasiliensis* infection, NLRP3 had an unexpected, and apparently inflammasome-independent, role in constraining the initial neutrophilia as well as the eosinophilia that results from the type 2 responses to the parasite and subsequent expulsion in the small intestine. These data suggest that the NLRP3 inflammasome has differential roles during innate and adaptive immunity and may be tissue-specific, acting differently in two very distinct helminth infection models.

We previously published that Ym1 was responsible for a more aggressive early neutrophilic response in the lung, which promotes *N. brasiliensis* larval killing but at the cost of increased host tissue damage (10). Ym1 can form crystals in Mφs (12) which may be a trigger for NLRP3 inflammasome assembly (35). Classically, neutrophilic influx, which can be induced by the presence of necrotic cells and DAMPs such as extracellular ATP during acute tissue injury, has been shown to be critically driven by NLRP3 (36). Additionally, NLRP3 inflammasome-dependent release of IL-1β is critical for neutrophil anti-bacterial responses to *Streptococcus pneumoniae* infection in the lungs (37). We therefore hypothesized that Ym1 acted via NLRP3, and NLRP3 deficiency would reverse Ym1-mediated neutrophil recruitment. However, our data counterintuitively showed that during acute lung injury with *N. brasiliensis* infection, NLRP3 restricted neutrophilia. In addition, we saw no evidence for inflammasome activation during innate responses and neither MCC950-mediated inflammasome inhibition, nor Caspase 1/11 deficiency, replicated NLRP3-deficiency during *N. brasiliensis* infection. It is therefore possible that NLRP3 has distinct roles regulating neutrophil recruitment that can either be dependent or independent of inflammasome activation. Our data are consistent with another report in which NLRP3, but not caspase-1/11 or ASC, limited early neutrophil influx during lung bacterial infection with *Francisella tularensis* (24). CXCL3, a neutrophil chemotactic factor that can mobilise neutrophils from the BM (38), was increased in the lungs of *N. brasiliensis* infected NLRP3-deficient mice, suggesting a potential mechanism for NLRP3-dependent regulation of neutrophil recruitment. However, the increased *Cxcl3* expression we observed may just reflect increased neutrophil numbers as recruited neutrophils can express *Cxcl3* (21).

In line with previous studies (5, 10), the enhanced neutrophilia we observed in the absence of NLRP3 resulted in increased lung damage and impaired repair mechanisms. The initiation of repair responses in the lung following injury due to *N. brasiliensis* migration is dependent on IL-4Rα signalling (5). Our transcriptional analysis revealed enhanced alternative Mφ activation and other type 2 markers such as *Il4* and *Arg1* in infected *Nlrp3*^−/−^ mice relative to WT controls. This correlated with enhanced *Ccl24* (Eotaxin-2) expression and increased eosinophilia in infected *Nlrp3*^−/−^ mice. Further, *Mmp13* expression was also elevated in infected *Nlrp3*^−/−^ mice. As MMP13 has been shown to be regulated by IL-4Rα during *N. brasiliensis* infection (5), the data further highlights possible type 2-dependent defects in tissue repair processes in the absence of NLRP3. Thus, in our model NLRP3 limits the host-protective type 2 response and thus in its absence, one might expect enhanced repair. However, paradoxically there was persistent damage in the gene deficient mice, suggesting NLRP3 is required for the timely resolution of the early inflammatory response. It is possible that type 2 responses are exaggerated in the absence of NLRP3 due to a compensatory need to repair the damaged tissues. Our data may also fit the hypothesis that early neutrophil responses are required to establish a type 2 response (39) and because we saw enhanced neutrophilia in NLRP3-deficient mice, the magnitude of the resulting type 2 response could be exacerbated.

These findings highlight a complex relationship between NLRP3 and IL-4. It has been previously reported that IL-4 signalling negatively regulates NLRP3 by suppressing inflammasome assembly and caspase-1 activity as well as the subcellular localization of NLRP3 (40). Additionally, in CD4^+^ T cells it has been shown that NLRP3 can act as a transcription factor that supports differentiation to the Th2 cell lineage in an inflammasome-independent manner (23). Our data suggest an additional mechanism by which NLRP3 can negatively regulate Mφ IL-4 responsiveness and controls expression of alternative activation markers during helminth infection in the lungs, independent of the inflammasome. Whether NLRP3 is acting as a transcription factor in Mφs similar to its action in Th2 cells and how mechanistically, NLRP3 regulates IL-4 signalling, will be the subject of future work. It will also be important to clarify how inflammasome-dependent and -independent roles for NLRP3 relate to innate versus adaptive immune responses within type 2 disease contexts.

## Materials and Methods

### Mice and ethics statements

*Nlrp3*^tm1Vmd^ (RRID:MGI:5468973) (41) (University of Manchester), B6-*Nlrp3*^tm1Tsc^/Siec (RRID:IMSR_MUGEN:M153001) (James Cook University (JCU)) and *Casp1/11*^−/−^ (RRID:IMSR_JAX:016621) (JCU) mice were maintained on a C57BL/6J (RRID:IMSR_JAX:000664) background and bred in-house at the University of Manchester and JCU. All experiments were carried out in accordance with the United Kingdom Animals (Scientific Procedures) Act 1986 and under a Project License (70/8548) granted by the Home Office and approved by local Animal Ethics Review Group at the University of Manchester. Animal experimental protocols in Australia were approved by the JCU Animal Ethics Committee (A2213).

### *N. brasiliensis* infection

*N. brasiliensis* worms were propagated as previously described (42). Infective L3 larvae were isolated and washed with sterile PBS and counted using a dissecting microscope. Mice were injected with 250 or 500 L3s subcutaneously. Upon culling the mice by pentobarbitone overdose i.p., BAL was performed with 10% FBS in PBS and lung lobes were collected. Lobes were either stored in RNAlater (Thermofisher), fixed in 10% formalin for histology or digested with Liberase TL (Roche). For lung-stage L4 counts, on days 1-2 post-infection, lungs were minced and incubated in PBS for 3 hours at 37 °C. Emergent larvae were counted using a dissecting microscope. For small intestinal worm burdens, worms were counted using a dissecting microscope following incubation at 37 °C. Fecal parasite eggs were enumerated from one fecal pellet/animal collected 6 days post-infection using a Whitlock paracytometer.

### NLRP3 inhibition

One day prior to *N. brasiliensis* infection, mice were treated with either PBS vehicle or 10 mg/kg MCC950 (Sigma) by intraperitoneal injection in a 100 μl volume, a dose which has been previously shown to inhibit inflammasome activity (43). Injections were repeated daily up until the experimental endpoint.

### IL-4 complex injection and *in vitro* IL-4 stimulation

For *in vivo* IL-4 stimulation, mice were injected intraperitoneally with 1.5 μg recombinant IL-4 complexed with 7.5 μg anti-IL-4. After 24 hours, lavage was performed to isolate peritoneal exudate cells containing predominantly Mφs which were analysed by flow cytometry. For *in vitro* stimulation, day 6 cultured BM-derived Mφs (differentiated with L929-conditioned media) were treated with 20 ng/ml IL-4 and analysed by flow cytometry the following day.

### *Ex vivo* caspase-1 activation assay

BAL cells were collected from mice at day 2 post-infection with *N. brasiliensis*. Cells were then stained for inflammasome activation using a caspase-1 FAM-YVAD-FMK FLICA probe kit (Bio-Rad). Cells were then processed for analysis by flow cytometry.

### Flow cytometry

Single cell suspensions were washed in PBS and Live/Dead staining (Thermofisher) was performed. Samples were Fc-blocked using α-CD16/32 (2.4G2, RRID:AB_394656) (BD Biosciences) and mouse serum (BioRad). Blocking and subsequent surface staining was performed using PBS containing 2 mM EDTA, 2% FBS, and 0.05% NaN_3_. Antibodies used for staining are listed in **Table 1**. Following surface staining, cells were incubated with IC fixation buffer (ThermoFisher) prior to permeabilization for intracellular staining. For secondary detection of Ym1 and RELM-α, Zenon goat (RRID:AB_2753200) and rabbit (RRID:AB_2572214) antibody labels (ThermoFisher) were used. For Ym1, RELM-α and pro-IL-1β intracellular staining, cells were directly stained without stimulation or protein transport inhibition. For cell quantification, some samples were spiked with 10 μm polystyrene beads (Sigma-Aldrich) prior to acquisition. Data were acquired on a BD LSRFortessa flow cytometer and analysed using FlowJo v10 software.

**Table 1.**
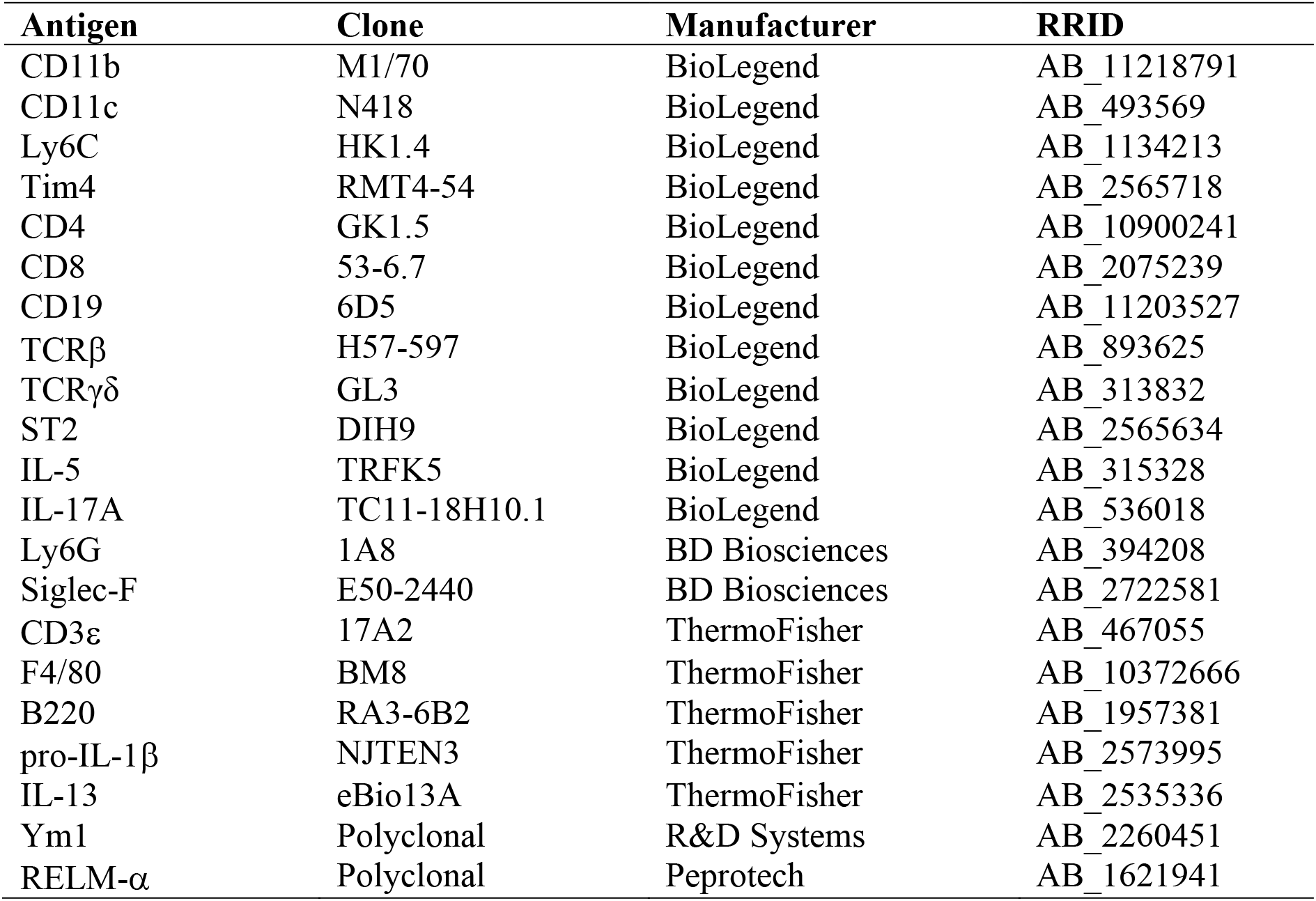
List of flow cytometry antibodies used.

### ELISA

BAL supernatants were analysed for Ym1 using commercially available ELISA kits (R&D systems and Peprotech, respectively). Lung homogenates were assayed for IL-4 using standard sandwich ELISA protocols (eBioscience). Analytes were detected using horse radish peroxidase-conjugated streptavidin and TMB substrate (BioLegend) and stopped with 1 M HCl. Final absorbance at 450 nm was measured using a VersaMax microplate reader (Molecular Devices).

### RNA extraction and quantitative real-time PCR

Tissue samples stored in RNAlater (Thermo Fisher Scientific) were processed for RNA extraction using a TissueLyser II and QIAzol reagent (Qiagen). Isolated RNA was quantified using a Qubit fluorimeter and RNA BR kit (Qiagen). cDNA was synthesized using Tetro reverse transcription kit (Bioline) and oligo dT 15-mers (Integrated DNA Technologies). Quantitative real-time PCR was performed using SYBR green mix (Agilent Technologies) and a LightCycler 480 II (Roche). A list of primer sequences used are shown in **Table 2**.

**Table 2.**
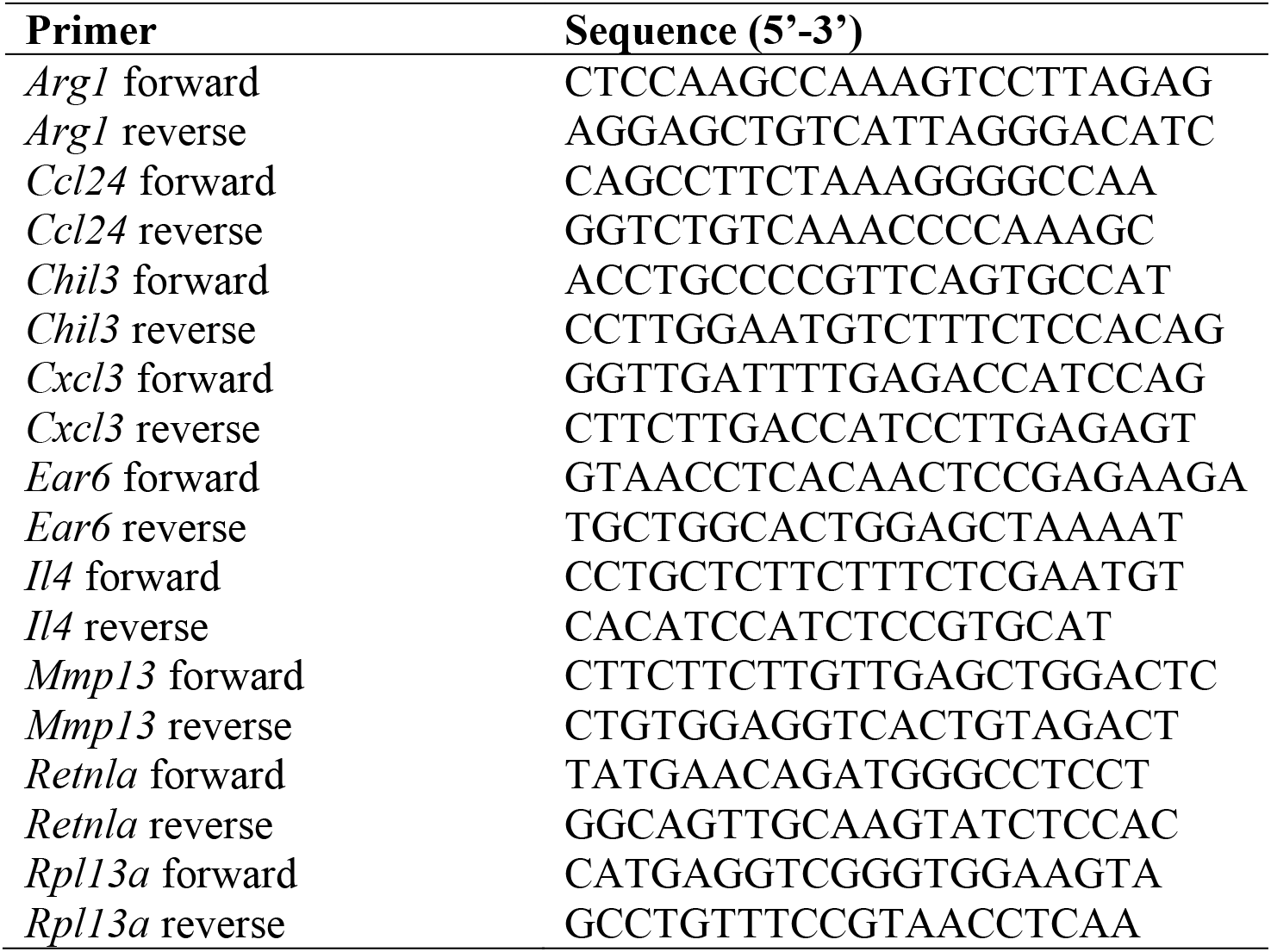
List of primer sequences used.

### Transcriptional profiling

Quality control was performed on RNA samples with an Agilent 2200 TapeStation system prior to downstream analyses. Samples were diluted and 100 ng of RNA was processed for running on a Nanostring nCounter® FLEX system using the Myeloid Innate Immunity v2 panel. Please note that this panel does not distinguish *Chil3* from *Chil4*. Raw counts were normalized to internal spike in controls and the expression of 13 stable housekeeping genes, as determined by geNorm algorithm within the nSolver Advanced Analysis tool. Subsequent analyses were performed in R (version 3.5.3). After normalization, transcripts with >15 counts were considered to be expressed and were log_2_ transformed. Linear modelling using the limma R package (44) was used to calculate differential gene expression. All expressed genes were used for principal components analysis (PCA). Unsupervised hierarchical clustering was performed on significantly differentially expressed genes between *N. brasiliensis*-infected WT and *Nlrp3*^−/−^ mice the using the linkage method complete and Euclidian distances.

### Histology and fractal analysis

Proximal small intestine was fixed in 4% formaldehyde and embedded in paraffin, and 5 μM sections were stained with PAS/Alcian Blue stains using the standard protocol of an institutional histology service provider (QIMR Berghofer Medical Research Institute). Goblet cells were quantified by counting PAS^+^ cells in 10 randomly selected villi units and averaged for each individual mouse. Whole left lung lobes were paraffin embedded and 5 μm sections were prepared for haematoxylin/eosin staining. Slides were imaged using an Olympus slide scanner and high-resolution image files were exported using Pannoramic Viewer software (3DHISTECH). The images were then processed in a KNIME software workflow to obtain 50 random regions of interests (ROIs) across the whole lung section. ROIs that contained lobe boundaries or extensive artefacts were excluded from the analysis. The ROIs were then converted to binary images and lacunarity (Λ) was quantified using the FracLac plugin for ImageJ (default settings). The Λ values of all the ROIs were averaged to obtain estimates for the entire lobe.

### Statistical analyses

Graphpad Prism 7 software was used for all statistical analyses. Data were assessed to be normally distributed by the D’Agostino-Pearson omnibus normality test. Differences between experimental groups were assessed by ANOVA (for normally distributed data) followed by Tukey-Kramer *post hoc* multiple comparisons test or an unpaired two-tailed Student’s *t* test. In cases where data were not normally distributed, a Kruskal-Wallis test was used. For gene expression data, values were log_2_ transformed to achieve normal distribution. Comparisons with a *P* value of less than 0.05 were considered to be statistically significant.

## Acknowledgements

This work was supported by the Wellcome Trust (106898/A/15/Z to JEA), the Medical Research Council UK (MR/K01207X/1 and MR/J001929/1 to JEA; MR/N003586/1 to DB), the Australian National Health and Medical Research Council (NHMRC grants 1117504 and 1132975 to AL), the Queensland Department of Science, Information Technology and Innovation (to PRG) and the Iraqi Cultural Attache in Australia (to RA). TES was supported by Medical Research Foundation UK joint funding with Asthma UK (MRFAUK-2015-302). We thank Vishva Dixit (Genentech) for the *Nlrp3*^tm1Vmd^ mice. Thanks to Seth Masters (WEHI, Australia) for kindly providing B6-*Nlrp3*^tm1Tsc^ /Siec and Caspase1/11 double-deficient mice. We thank the Flow Cytometry, Bioimaging, Genomic Technologies, and Biological Services core facilities at the University of Manchester.

## Author Contributions

ALC, RA, AK, AABR, TES, PRG and JEA designed the research plan. ALC, RA, ZA, JA, JP, MMC, BHKC and RME performed experiments, analysed data and revised the manuscript. LAD, AABR, AK, DB, AL and TES provided materials, intellectual input and revised the manuscript. ALC, PRG, and JEA wrote the manuscript. JEA, PRG and AL supervised the work and provided funding.

## References

1. Martinez, F. O. 2008. Macrophage activation and polarization. Front. Biosci. 13: 453.

2. Sutherland, T. E., D. Rückerl, N. Logan, S. Duncan, T. A. Wynn, and J. E. Allen. 2018. Ym1 induces RELMα and rescues IL-4Rα deficiency in lung repair during nematode infection. PLOS Pathog. 14: e1007423.

3. Kreider, T., R. M. Anthony, J. F. Urban, W. C. Gause, and W. C. Gause. 2007. Alternatively activated macrophages in helminth infections. Curr. Opin. Immunol. 19: 448–453.

4. Knipper, J. A., S. Willenborg, J. Brinckmann, W. Bloch, T. Maaß, R. Wagener, T. Krieg, T. Sutherland, A. Munitz, M. E. Rothenberg, A. Niehoff, R. Richardson, M. Hammerschmidt, J. E. Allen, and S. A. Eming. 2015. Interleukin-4 receptor α signaling in myeloid cells controls collagen fibril assembly in skin repair. Immunity 43: 803–816.

5. Chen, F., Z. Liu, W. Wu, C. Rozo, S. Bowdridge, A. Millman, N. Van Rooijen, J. F. Urban, T. A. Wynn, W. C. Gause, and W. C. Gause. 2012. An essential role for TH2-type responses in limiting acute tissue damage during experimental helminth infection. Nat. Med. 18: 260–266.

6. Harvie, M., M. Camberis, S.-C. Tang, B. Delahunt, W. Paul, and G. Le Gros. 2010. The lung is an important site for priming CD4 T-cell-mediated protective immunity against gastrointestinal helminth parasites. Infect. Immun. 78: 3753–62.

7. Bouchery, T., R. Kyle, M. Camberis, A. Shepherd, K. Filbey, A. Smith, M. Harvie, G. Painter, K. Johnston, P. Ferguson, R. Jain, B. Roediger, B. Delahunt, W. Weninger, E. Forbes-Blom, and G. Le Gros. 2015. ILC2s and T cells cooperate to ensure maintenance of M2 macrophages for lung immunity against hookworms. Nat. Commun. 6: 6970.

8. Obata-Ninomiya, K., K. Ishiwata, H. Nakano, Y. Endo, T. Ichikawa, A. Onodera, K. Hirahara, Y. Okamoto, H. Kanuka, and T. Nakayama. 2018. CXCR6+ ST2+ memory T helper 2 cells induced the expression of major basic protein in eosinophils to reduce the fecundity of helminth. Proc. Natl. Acad. Sci. 115: E9849–E9858.

9. Chen, F., W. Wu, A. Millman, J. F. Craft, E. Chen, N. Patel, J. L. Boucher, J. F. Urban, C. C. Kim, and W. C. Gause. 2014. Neutrophils prime a long-lived effector macrophage phenotype that mediates accelerated helminth expulsion. Nat. Immunol. 15: 938–946.

10. Sutherland, T. E., N. Logan, D. Rückerl, A. A. Humbles, S. M. Allan, V. Papayannopoulos, B. Stockinger, R. M. Maizels, and J. E. Allen. 2014. Chitinase-like roteins promote IL-17-mediated neutrophilia in a tradeoff between nematode killing and host damage. Nat. Immunol. 15: 1116–1125.

11. Muallem, G., and C. A. Hunter. 2014. ParadYm shift: Ym1 and Ym2 as innate immunological regulators of IL-17. Nat. Immunol. 15: 1099–1100.

12. Liu, Q., L. I. Cheng, L. Yi, N. Zhu, A. Wood, C. M. Changpriroa, J. M. Ward, and S. H. Jackson. 2009. p47phox deficiency induces macrophage dysfunction resulting in progressive crystalline macrophage pneumonia. Am. J. Pathol. 174: 153–163.

13. Alhallaf, R., Z. Agha, C. M. Miller, A. A. B. Robertson, J. Sotillo, J. Croese, M. A. Cooper, S. L. Masters, A. Kupz, N. C. Smith, A. Loukas, and P. R. Giacomin. 2018. The NLRP3 inflammasome suppresses protective immunity to gastrointestinal helminth infection. Cell Rep. 23: 1085–1098.

14. Chen, F., Z. Liu, W. Wu, C. Rozo, S. Bowdridge, A. Millman, N. Van Rooijen, J. F. Urban, T. A. Wynn, W. C. Gause, and W. C. Gause. 2012. An essential role for TH2-type responses in limiting acute tissue damage during experimental helminth infection. Nat. Med. 18: 260–266.

15. Shimokawa, C., T. Kanaya, M. Hachisuka, K. Ishiwata, H. Hisaeda, Y. Kurashima, H. Kiyono, T. Yoshimoto, T. Kaisho, and H. Ohno. 2017. Mast cells are crucial for induction of group 2 innate lymphoid cells and clearance of helminth infections. Immunity 46: 863–874.e4.

16. Gause, W. C., T. A. Wynn, and J. E. Allen. 2013. Type 2 immunity and wound healing: evolutionary refinement of adaptive immunity by helminths. Nat. Rev. Immunol. 13: 607–614.

17. Reece, J. J., M. C. Siracusa, and A. L. Scott. 2006. Innate immune responses to lung-stage helminth infection induce alternatively activated alveolar macrophages. Infect. Immun. 74: 4970–4981.

18. Marsland, B. J., M. Kurrer, R. Reissmann, N. L. Harris, and M. Kopf. 2008. Nippostrongylus brasiliensis infection leads to the development of emphysema associated with the induction of alternatively activated macrophages. Eur. J. Immunol. 38: 479–488.

19. Porzionato, A., D. Guidolin, V. Macchi, G. Sarasin, D. Grisafi, C. Tortorella, A. Dedja, P. Zaramella, and R. De Caro. 2016. Fractal analysis of alveolarization in hyperoxia-induced rat models of bronchopulmonary dysplasia. Am. J. Physiol. Cell. Mol. Physiol. 310: L680–L688.

20. Benjamini, Y., and Y. Hochberg. 1995. Controlling the false discovery rate: a practical and powerful approach to multiple testing. J. R. Stat. Soc. Ser. B 57: 289–300.

21. Shay, T., and J. Kang. 2013. Immunological Genome Project and systems immunology. Trends Immunol. 34: 602–609.

22. Gundra, U. M., N. M. Girgis, D. Ruckerl, S. Jenkins, L. N. Ward, Z. D. Kurtz, K. E. Wiens, M. S. Tang, U. Basu-Roy, A. Mansukhani, J. E. Allen, and P. Loke. 2014. Alternatively activated macrophages derived from monocytes and tissue macrophages are phenotypically and functionally distinct. Blood 123: e110–22.

23. Bruchard, M., C. Rebé, V. Derangère, D. Togbé, B. Ryffel, R. Boidot, E. Humblin, A. Hamman, F. Chalmin, H. Berger, A. Chevriaux, E. Limagne, L. Apetoh, F. Végran, and F. Ghiringhelli. 2015. The receptor NLRP3 is a transcriptional regulator of TH2 differentiation. Nat. Immunol. 16: 859–870.

24. Periasamy, S., H. T. Le, E. B. Duffy, H. Chin, and J. A. Harton. 2016. Inflammasome-independent NLRP3 restriction of a protective early neutrophil response to pulmonary tularemia. PLOS Pathog. 12: e1006059.

25. Boucher, D., M. Monteleone, R. C. Coll, K. W. Chen, C. M. Ross, J. L. Teo, G. A. Gomez, C. L. Holley, D. Bierschenk, K. J. Stacey, A. S. Yap, J. S. Bezbradica, and K. Schroder. 2018. Caspase-1 self-cleavage is an intrinsic mechanism to terminate inflammasome activity. J. Exp. Med. 215: 827–840.

26. Schroder, K., and J. Tschopp. 2010. The inflammasomes. Cell 140: 821–832.

27. Madouri, F., N. Guillou, L. Fauconnier, T. Marchiol, N. Rouxel, P. Chenuet, A. Ledru, L. Apetoh, F. Ghiringhelli, M. Chamaillard, S. G. Zheng, F. Trovero, V. F. J. Quesniaux, B. Ryffel, and D. Togbe. 2015. Caspase-1 activation by NLRP3 inflammasome dampens IL-33-dependent house dust mite-induced allergic lung inflammation. J. Mol. Cell Biol. 7: 351–365.

28. Coll, R. C., A. A. B. Robertson, J. J. Chae, S. C. Higgins, R. Muñoz-Planillo, M. C. Inserra, I. Vetter, L. S. Dungan, B. G. Monks, A. Stutz, D. E. Croker, M. S. Butler, M. Haneklaus, C. E. Sutton, G. Núñez, E. Latz, D. L. Kastner, K. H. G. Mills, S. L. Masters, K. Schroder, M. A. Cooper, and L. A. J. O’Neill. 2015. A small-molecule inhibitor of the NLRP3 inflammasome for the treatment of inflammatory diseases. Nat. Med. 21: 248–255.

29. Kool, M., V. Pétrilli, T. De Smedt, A. Rolaz, H. Hammad, M. van Nimwegen, I. M. Bergen, R. Castillo, B. N. Lambrecht, and J. Tschopp. 2008. Alum adjuvant stimulates inflammatory dendritic cells through activation of the NALP3 inflammasome. J. Immunol. 181: 3755–3759.

30. Franchi, L., and G. Núñez. 2008. The NLRP3 inflammasome is critical for aluminium hydroxide-mediated IL-1beta secretion but dispensable for adjuvant activity. Eur. J. Immunol. 38: 2085–2089.

31. Besnard, A.-G., N. Guillou, J. Tschopp, F. Erard, I. Couillin, Y. Iwakura, V. Quesniaux, B. Ryffel, and D. Togbe. 2011. NLRP3 inflammasome is required in murine asthma in the absence of aluminum adjuvant. Allergy 66: 1047–1057.

32. Primiano, M. J., B. A. Lefker, M. R. Bowman, A. G. Bree, C. Hubeau, P. D. Bonin, M. Mangan, K. Dower, B. G. Monks, L. Cushing, S. Wang, J. Guzova, A. Jiao, L.-L. Lin, E. Latz, D. Hepworth, and J. P. Hall. 2016. Efficacy and pharmacology of the NLRP3 inflammasome inhibitor CP-456,773 (CRID3) in murine models of dermal and pulmonary inflammation. J. Immunol. 197: 2421–2433.

33. Kim, R. Y., J. W. Pinkerton, A. T. Essilfie, A. A. B. Robertson, K. J. Baines, A. C. Brown, J. R. Mayall, M. K. Ali, M. R. Starkey, N. G. Hansbro, J. A. Hirota, L. G. Wood, J. L. Simpson, D. A. Knight, P. A. Wark, P. G. Gibson, L. A. J. O’Neill, M. A. Cooper, J. C. Horvat, and P. M. Hansbro. 2017. Role for NLRP3 inflammasome-mediated, IL-1β-dependent responses in severe, steroid-resistant asthma. Am. J. Respir. Crit. Care Med. 196: 283–297.

34. Huang, H., J.-Y. Hong, Y.-J. Wu, E.-Y. Wang, Z.-Q. Liu, B.-H. Cheng, L. Mei, Z.-G. Liu, P.-C. Yang, and P.-Y. Zheng. 2018. Vitamin D receptor interacts with NLRP3 to restrict the allergic response. Clin. Exp. Immunol. 194: 17–26.

35. Persson, E. K., K. Verstraete, I. Heyndrickx, E. Gevaert, H. Aegerter, J.-M. Percier, K. Deswarte, K. H. Verschueren, A. Dansercoer, D. Gras, P. Chanez, C. Bachert, A. Goncalves, H. van Gorp, H. de Haard, C. Blanchetot, M. Saunders, H. Hammad, S. N. Savvides, and B. N. Lambrecht. 2019. Protein crystallization promotes type 2 immunity and is reversible by antibody treatment. Science 364: eaaw4295.

36. Iyer, S. S., W. P. Pulskens, J. J. Sadler, L. M. Butter, G. J. Teske, T. K. Ulland, S. C. Eisenbarth, S. Florquin, R. A. Flavell, J. C. Leemans, and F. S. Sutterwala. 2009. Necrotic cells trigger a sterile inflammatory response through the NLRP3 inflammasome. Proc. Natl. Acad. Sci. 106: 20388–20393.

37. Karmakar, M., M. Katsnelson, H. A. Malak, N. G. Greene, S. J. Howell, A. G. Hise, A. Camilli, A. Kadioglu, G. R. Dubyak, and E. Pearlman. 2015. Neutrophil IL-1β processing induced by pneumolysin is mediated by the NLRP3/ASC inflammasome and caspase-1 activation and is dependent on K + efflux. J. Immunol. 194: 1763–1775.

38. Burdon, P. C. E., C. Martin, and S. M. Rankin. 2005. The CXC chemokine MIP-2 stimulates neutrophil mobilization from the rat bone marrow in a CD49d-dependent manner. Blood 105: 2543–2548.

39. Tacchini-Cottier, F., C. Zweifel, Y. Belkaid, C. Mukankundiye, M. Vasei, P. Launois, G. Milon, and J. A. Louis. 2000. An immunomodulatory function for neutrophils during the induction of a CD4+ Th2 response in BALB/c mice infected with Leishmania major. J. Immunol. 165: 2628–2636.

40. Hwang, I., J. Yang, S. Hong, E. Ju Lee, S.-H. Lee, T. Fernandes-Alnemri, E. S. Alnemri, and J.-W. Yu. 2015. Non-transcriptional regulation of NLRP3 inflammasome signaling by IL-4. Immunol. Cell Biol. 93: 591–599.

41. Mariathasan, S., D. S. Weiss, K. Newton, J. McBride, K. O’Rourke, M. Roose-Girma, W. P. Lee, Y. Weinrauch, D. M. Monack, and V. M. Dixit. 2006. Cryopyrin activates the inflammasome in response to toxins and ATP. Nature 440: 228–232.

42. Lawrence, R. A., C. A. Gray, J. Osborne, and R. M. Maizels. 1996. Nippostrongylus brasiliensis: cytokine responses and nematode expulsion in normal and IL-4-deficient mice. Exp. Parasitol. 84: 65–73.

43. Krishnan, S. M., Y. H. Ling, B. M. Huuskes, D. M. Ferens, N. Saini, C. T. Chan, H. Diep, M. M. Kett, C. S. Samuel, B. K. Kemp-Harper, A. A. B. Robertson, M. A. Cooper, K. Peter, E. Latz, A. S. Mansell, C. G. Sobey, G. R. Drummond, and A. Vinh. 2019. Pharmacological inhibition of the NLRP3 inflammasome reduces blood pressure, renal damage, and dysfunction in salt-sensitive hypertension. Cardiovasc. Res. 115: 776–787.

44. Ritchie, M. E., B. Phipson, D. Wu, Y. Hu, C. W. Law, W. Shi, and G. K. Smyth. 2015. limma powers differential expression analyses for RNA-sequencing and microarray studies. Nucleic Acids Res. 43: e47–e47.

