## supplemental Data for "Inflammasome-independent role for NLRP3 in controlling innate anti-helminth immunity and tissue repair in the lung"

### Supplemental Material

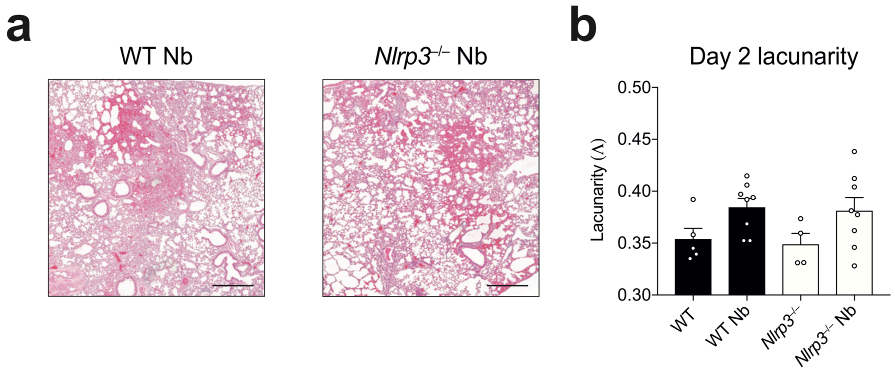


**Figure S1** Lung histopathology of WT and *Nlrp3*$^{-/-}$ mice on Day 2 post-infection with *N. brasiliensis*. (**a**) Haematoxylin/eosin staining was performed on lung sections and imaged followed by (**b**) quantification of lacunarity (Λ) to assess lung damage. Data are pooled (**b**; mean ± s.e.m.) from 2 individual experiments with 3-4 mice per group (per experiment).

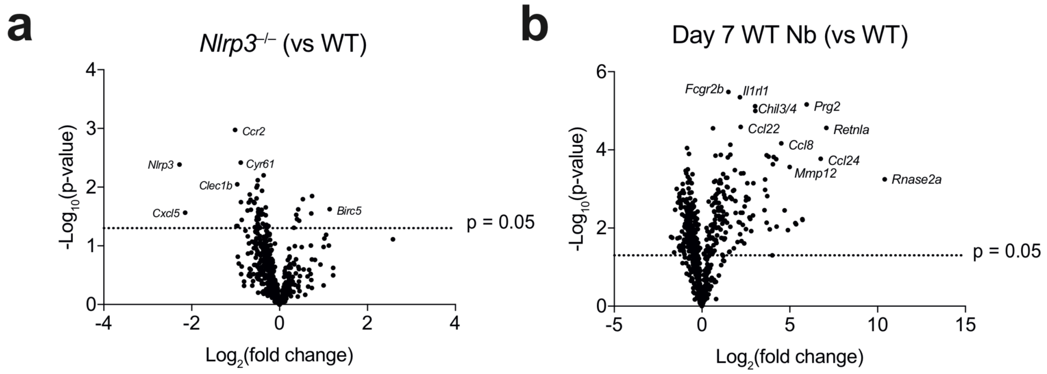


**Figure S2** Differential gene expression in naïve *Nlrp3*$^{-/-}$ mice and most upregulated genes during *N. brasiliensis* infection. (**a**) Whole lung RNA was analysed by NanoString and volcano plots were generated to assess differential gene expression in naïve *Nlrp3*$^{-/-}$ mice compared to naïve WT mice. (**b**) Differential gene expression in WT mice infected with *N. brasiliensis* on day 7 post-infection compared to WT naïve mice. Data were from a single Nanostring run (n=2-4 per group).

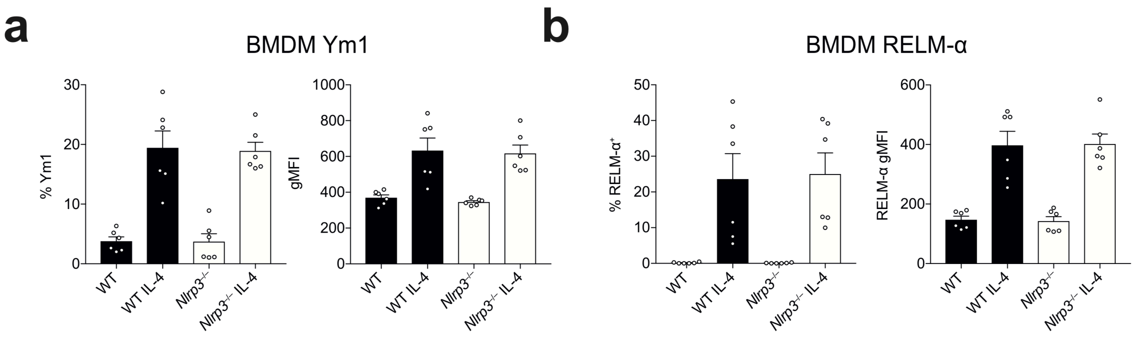

**Figure S3** IL-4 stimulation of BMDM cultures. BMDMs were generated from WT and *Nlrp3*$^{-/-}$ mice and stimulated for 24 hours with IL-4. Frequencies and gMFI are shown for (**a**) Ym1 and (**b**) RELM-α for CD11b$^{+}$ BMDMs analysed by flow cytometry. Data are pooled from 2 independent experiments (mean ± s.e.m.) from 2 individual experiments with 3-4 mice per group (per experiment).

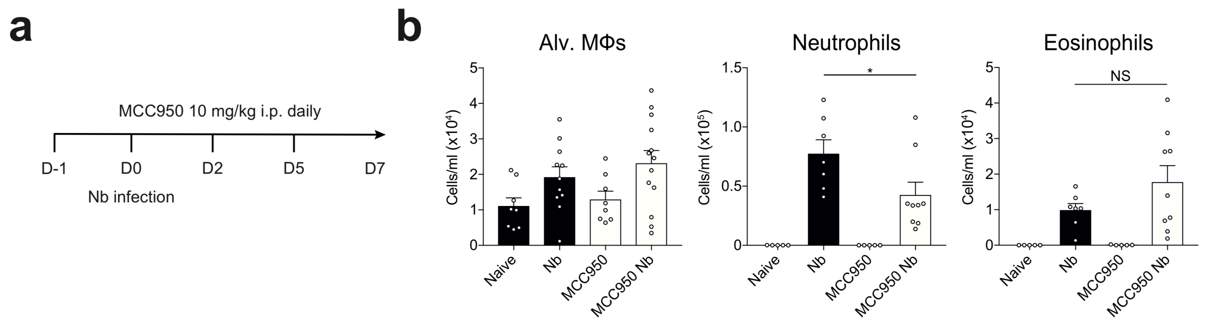


**Figure S4** MCC950 treatment does not phenocopy *Nlrp3*$^{-/-}$ mice during *N. brasiliensis* infection. (**a**) WT mice were treated with MCC950 (time course shown) and (**b**) absolute numbers of BAL alveolar Mφs, neutrophils, and eosinophils on day 5 post-infection were determined by flow cytometry. Data are pooled (**b**; mean ± s.e.m.) from 3 individual experiments with 3-5 mice per group (per experiment). *$P$<0.05 (one-way ANOVA and Tukey-Kramer *post hoc* test).
